# Kmer2SNP: reference-free SNP calling from raw reads based on matching

**DOI:** 10.1101/2020.05.17.100305

**Authors:** Yanbo Li, Yu Lin

**Affiliations:** Research School of Computer Science, Australian National University

**Keywords:** SNP calling, k-mer frequency distribution, reference free, maximum weight matching

## Abstract

The development of DNA sequencing technologies provides the opportunity to call heterozygous SNPs for each individual. SNP calling is a fundamental problem of genetic analysis and has many applications, such as gene-disease diagnosis, drug design, and ancestry inference. Reference-based SNP calling approaches generate highly accurate results, but they face serious limitations especially when high-quality reference genomes are not available for many species. Although reference-free approaches have the potential to call SNPs without using the reference genome, they have not been widely applied on large and complex genomes because existing approaches suffer from low recall/precision or high runtime.

We develop a reference-free algorithm Kmer2SNP to call SNP directly from raw reads. Kmer2SNP first computes the k-mer frequency distribution from reads and identifies potential heterozygous k-mers which only appear in one haplotype. Kmer2SNP then constructs a graph by choosing these heterozygous k-mers as vertices and connecting edges between pairs of heterozygous k-mers that might correspond to SNPs. Kmer2SNP further assigns a weight to each edge using overlapping information between heterozygous k-mers, computes a maximum weight matching and finally outputs SNPs as edges between k-mer pairs in the matching.

We benchmark Kmer2SNP against reference-free methods including hybrid (assembly-based) and assembly-free methods on both simulated and real datasets. Experimental results show that Kmer2SNP achieves better SNP calling quality while being an order of magnitude faster than the state-of-the-art methods. Kmer2SNP shows the potential of calling SNPs only using k-mers from raw reads without assembly. The source code is freely available at https://github.com/yanboANU/Kmer2SNP.

## 1 Background

Whole-genome sequencing provides reads originated from two homologous sets of chromosomes (*i.e.*, haplotypes) from a single individual, thus makes it amenable to call heterozygous SNPs. Thanks to its high throughput, moderate cost and low error rate, next-generation sequencing (NGS) is gaining popularity in SNP calling with many applications in population genetics and biomedical research, such as gene-disease diagnosis, drug design and ancestry inference [31].

Most existing approaches for SNP calling rely on the alignment between raw reads and a reference genome. In general, the first step of those reference-based approaches is to align reads onto a reference genome, such as PolyScan[8], MAQ[18], SOAPsnp[19], GATK[9] and SAMtools[17]. However, aligning reads to a reference genome has some serious limitations. Firstly, the individual genome may contain regions that are absent (e.g., horizontal transfer elements) or divergent (e.g., HLA genes) from the reference genome. Therefore, short reads from these regions may not be able to correctly map to the reference genome. Secondly, short reads are prone to misalignment due to repeats in the reference genome, and the misalignment may affect the quality of SNP calling, especially in repetitive regions. Thirdly, a high-quality reference genome may not be available for read mapping and basepair-level inaccuracies in the reference genome may result in false positive SNPs.

There is a strong need to develop reference-free algorithms for SNP calling. Without the reference genome, such algorithms are limited to identify heterozygous SNPs (two different alleles on two haplotypes) rather than homozygous SNPs (the same allele on two haplotypes but differ from the reference). Moreover, the inferred SNPs cannot be assigned to a known genomic position as the reference genome is unknown. In hybrid approaches, raw reads are assembled into long contigs or scaffolds and SNPs can be identified by aligning raw reads to assembled contigs and assigned to positions on these contigs (instead of a reference genome). Such hybrid approaches suffer not only from misalignment and errors of raw reads but also from incompleteness and errors in assemblies [33]. Existing hybrid approaches are limited to call SNPs in organisms with a small genome size, and result in lower recall and precision rates comparing to the reference-based approaches [35]. Another type of reference-free algorithms call SNPs without explicitly assembling raw reads into contigs [35]. For example, de Bruijn graphs are built from raw reads and then SNPs are detected and represented as specific patterns *(e.g.,* bubble) in de Bruijn graphs [13,15,27,33,26]. However, detecting patterns becomes challenging when de Bruijn graphs get complicated due to various repeats in genomes and errors in raw reads. For example, bubbles may be incorporated into large bulges, resulting in low sensitivity in detecting SNPs. Moreover, EBWT2SNP [28] does not build a de Bruijn graph and uses extended Burrows-Wheeler Transform(eBWT) from reads to call SNPs as pairs of *k*-mers. However, the eBWT positional clustering may become ambiguous due to sequencing errors in raw reads or highly repetitive genomic regions, resulting in unstable SNP calling performance. Moreover, all the above reference-free algorithms are not very scalable because it is time-consuming to build either de Bruijn graphs or eBWT indices from all raw reads.

In this paper, we propose a reference-free approach, Kmer2SNP for SNP calling directly from raw reads. Our work has three contributions: (1) we show that the k-mer frequency distribution provides the power to detect heterozygous k-mers covering SNPs in an unknown reference genome; (2) we propose a graph model to model heterozygous k-mers and reduce the SNP calling problem to finding a maximum weight matching in the heterozygous k-mer graph; (3) experimental results show that Kmer2SNP achieves better quality while being an order of magnitude faster, compared to the state-of-the-art approaches for SNP calling without references.

## 2 Method

### 2.1 Preliminaries

In this section, we define some general terminology. Kmer2SNP is based on the k-mer analysis from raw reads. A k-mer is a subsequence of length k from either the reference genome or raw reads. Let *Σ* = {*A,C,G,T*} be the alphabet of nucleotides and *Σ^k^* represents the set of all k-mers. The *i^th^* nucleotide of a *k*-mer *x* is denoted as *x*[*i*], 1 ≤ *i* ≤ *k*. A segment of *k*-mer can be represented as *x*[*i,j*] = *x*[*i*]*x*[*i* + 1]… *x*[*j*], 1 ≤ *i* ≤ *j ≤ k*. A k-mer from raw reads is called *erroneous* if it does not appear in the reference genome. A k-mer from raw reads is called *genomic* if it appears in the reference genome. Genomic k-mers can be divided into two categories, *heterozygous* k-mers and *homozygous* k-mers. Heterozygous k-mers appear in only one of the two haplotypes while homozygous k-mers appear in both haplotypes. See Fig. 1 (A) for an example.

**Fig. 1.**
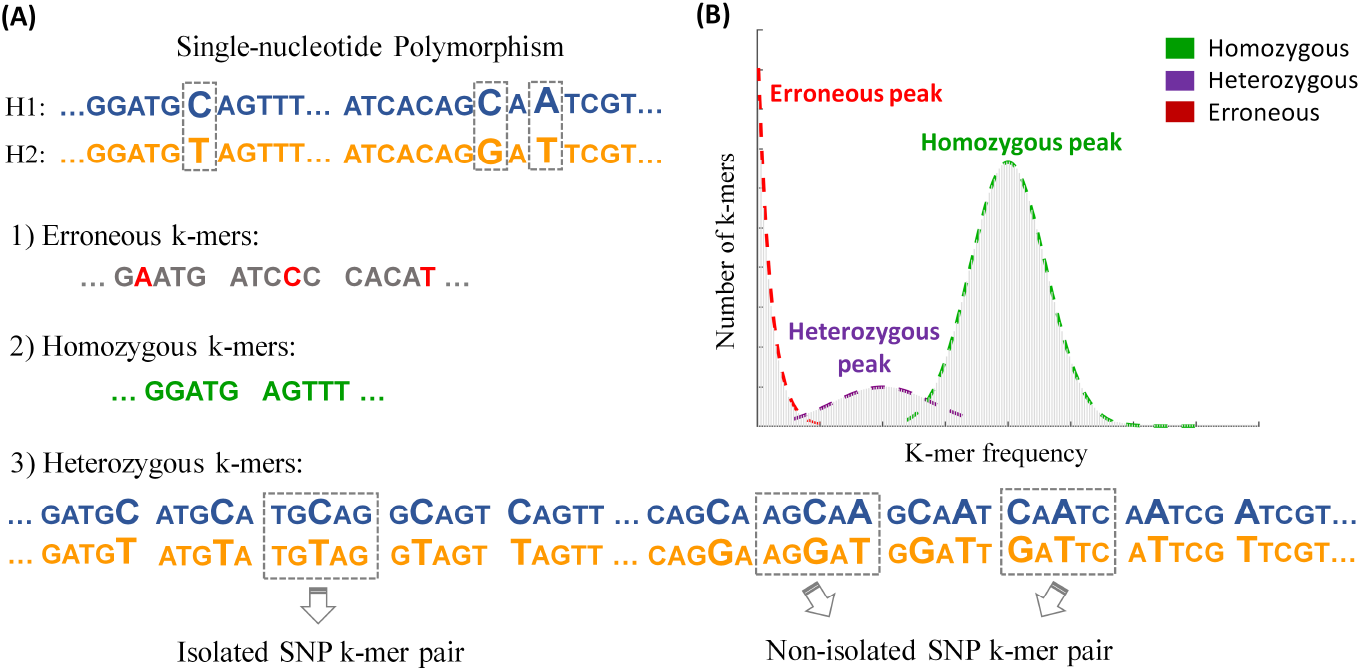
Erroneous, homozygous and heterozygous k-mers as well as isolated and non-isolated SNP k-mer pairs. (A) Example of erroneous, heterozygous and homozygous k-mers from raw reads of two haplotypes. Erroneous k-mers contain sequencing errors, heterozygous k-mers appear in only one haplotype and homozygous k-mers appear in both haplotypes. An isolated SNP corresponds to a pair of k-mers differ only at the middle position, i.e., an isolated SNP k-mer pair. A non-isolated SNP corresponds to two pairs of k-mers differ at least two positions (including the middle position), i.e., a non-isolated SNP k-mer pair. (B) K-mer frequency histogram built from raw reads. The x-axis represents k-mer frequency and y-axis represents the corresponding k-mer counts. Three peaks (from left to right) correspond to erroneous, heterozygous and homozygous k-mers, respectively.

We define the frequency of a k-mer as the number of times this k-mer appears in all raw reads and the frequency of k-mers (for sufficiently large k) can be used to distinguish erroneous, heterozygous and homozygous k-mers [32,34]. In an ideal case, erroneous k-mers have the lowest frequencies because sequencing errors are more or less random in raw reads. The frequencies of heterozygous k-mers are roughly half of the frequencies of homozygous k-mers as the former ones only appear in one of the two haplotypes. Fig. 1 (B) shows the k-mer frequency histogram where three peaks (from left to right), corresponding to erroneous, heterozygous and homozygous k-mers, respectively.

In the following, we focus on heterozygous k-mers in our work. A SNP can be represented by a pair of k-mers (from two haplotypes) that differ at the middle position. A SNP is called *isolated* if the above pair of k-mers differ only at the middle position and such a k-mer pair is called *isolated SNP* k-mer pair. A SNP is called *non-isolated* if the above pair of k-mers differ at least two positions (including the middle position) and such a k-mer pair is called *non-isolated SNP* k-mer pair. Fig. 1(A) shows examples of isolated SNP and non-isolated SNP along with their corresponding k-mer pairs.

### 2.2 Problem Formulation and Pipeline

Similar to the k-mer Hamming graph for error correction [22, 25], we introduce a graph model of heterozygous k-mers for SNP calling.

As shown in Fig. 2, Kmer2SNP takes raw reads as the input. The pipeline of Kmer2SNP starts from building a *heterozygous k-mer graph,* where vertices correspond to heterozygous k-mers and edges between a pair of heterozygous k-mers correspond to potential SNPs. Kmer2SNP further computes the weight for each edge using overlapping information between heterozygous k-mers. Kmer2SNP finally finds a maximum weight matching in the above graph and outputs the corresponding SNPs. We will explain each step in the following parts.

**Fig. 2.**
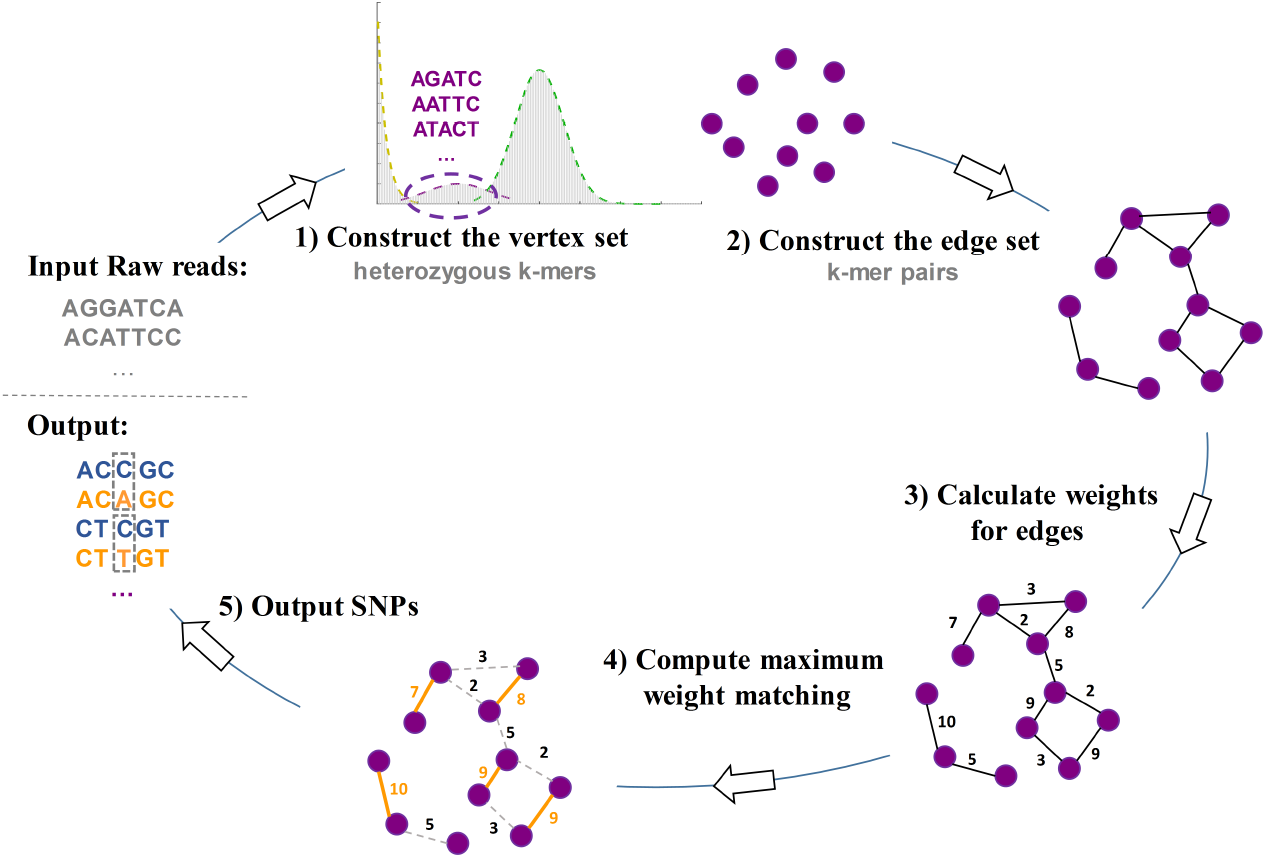
The pipeline of Kmer2SNP. Kmer2SNP takes raw reads as the input and starts from building a *heterozygous k-mer graph,* where vertices correspond to heterozygous k-mers and edges between a pair of heterozygous k-mers correspond to potential SNPs. Kmer2SNP further computes the weight for each edge using overlapping information between heterozygous k-mers. Kmer2SNP finally finds a maximum weight matching in the above graph and outputs the corresponding SNPs.

#### Step 1: Construct the vertex set – heterozygous k-mers

In the above heterozygous k-mer graph, each vertex is a heterozygous k-mer. Constructing the vertex set is to identify heterozygous k-mers. Kmer2SNP first counts the k-mer frequency from raw reads. K-mer counting is well-studied in computational genomics and there are many efficient tools available such as Jellyfish [21] and DSK [29]. Kmer2SNP uses DSK [29] to count k-mer frequencies from raw reads and derive a corresponding k-mer histogram file. We then use FindGSE [32] to find the frequency range of heterozygous k-mers. FindGSE fits the skew normal distributions to the above k-mer histogram and distinguishes peaks corresponding to erroneous, heterozygous and homozygous k-mers.

Using DSK and FindGSE, Kmer2SNP effectively retains most heterozygous k-mers and filters most erroneous k-mers and homozygous k-mers^1^. We use a simulated dataset from NA12878 Chromosome 22 (30x for each haplotype), HG-C_22_, as a running example. Even through only 9.8% of all k-mers (3,344,254 out of 34,011,172) in HG-C_22_ form the vertex set, they contain 99.34% heterozygous k-mers (1,399,216 out of 1,408,482). Kmer2SNP successfully filters 93.50% homozygous k-mers (27,826,482 out of 29,762,470) and 99.50% erroneous k-mers (1,794,489 out of 1,803,539). However, the vertex set still consists of 1,935,988 homozygous k-mers (57.89%) and 9,050 erroneous k-mers (0.27%), mixed with 1,399,216 heterozygous k-mers (41.84%). Kmer2SNP further uses k-mer pairing information to filter homozygous k-mers and erroneous k-mer in the next step.

#### Step 2: Construct the edge set – k-mers pairs

After constructing the vertex set, Kmer2SNP constructs edges between k-mer pairs that correspond to SNPs. As defined in Section 2.1, there are two types of k-mer pairs: isolated SNP k-mer pairs and non-isolated SNP k-mer pairs. Kmer2SNP thus introduces two types of edges, h1-edges and h2-edges to represent those k-mer pairs, respectively.

According to the definition of isolated SNP k-mer pairs, there is an *h*1-edge between two k-mers *x* and *x^ř^* if and only if 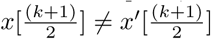 and *h*(*x,x*′) = 1, where *k* is odd and *h*() computes the hamming distance. KmerSNP does not calculate all-vs-all pairwise hamming distances between vertices due to the high running time. Instead, for each k-mer, Kmer2SNP removes its middle position and uses the other (k-1) positions as a key to compute its index in a hash table. Obviously, Kmer2SNP connects two k-mers *x* and *x*′ by an *h*1-edge if and only if they have the same index in the above hash table. Kmer2SNP also builds a similar hashing function to find non-isolated SNP k-mer pairs and construct corresponding *h*2-edges. Please refer to the Supplementary Section 1 for more details.

Following the same example dataset HG-C_22_ given in the Section 2.2 Step 1, the vertex set consists of 41.84% heterozygous k-mers, 57.89% homozygous k-mers and 0.27% erroneous k-mers. After constructing edges in the heterozygous k-mer graph, Kmer2SNP only retains connected components with at least 2 vertices and these connected components now consist of 99.36% heterozygous k-mers, 0.46% homozygous k-mers and 0.18% erroneous k-mers. Note that most homozygous k-mers at low coverage regions have been successfully filtered because they are unlikely to form any k-mer pairs. Although k-mers in connected components of size 2 (i.e., k-mer pairs) naturally correspond to potential SNPs, there are still a significant number of vertices locating in connected components with at least 3 vertices. If a k-mer is attached to multiple edges, how could we assign this k-mer to an edge that most likely corresponds to a real SNP? In the next step, we show how to compute a weight for each edge that indicates the likelihood of the corresponding SNP being true.

#### Step 3: Calculate weights for edges

In the heterozygous k-mer graph, each edge now corresponds to a potential SNP and the weight of an edge should indicate how likely this potential SNP is true. As shown Fig. 3 (a), an edge is constructed between an isolated SNP k-mer pair where these two k-mers only differ at the middle position. Note that are other heterozygous k-mers which also cover this isolated SNP (not at the middle position) and may provide support to this potential SNP. Therefore, Kmer2SNP calculates a weight for each edge based on the presence or absence overlapping heterozygous k-mers.

**Fig. 3.**
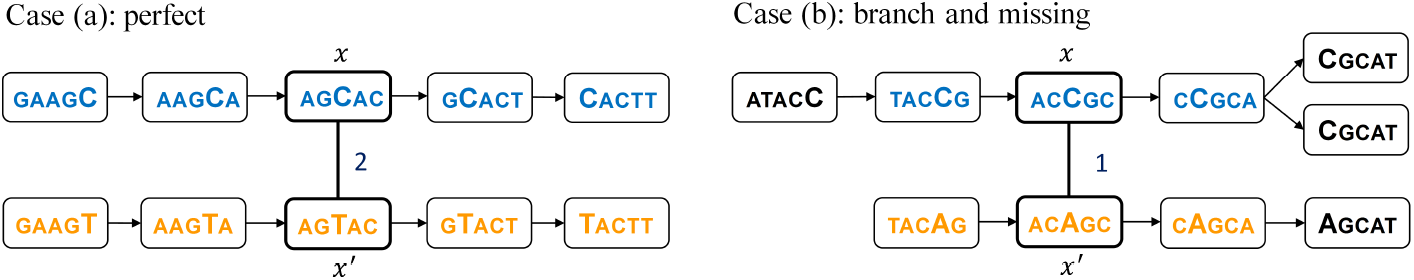
Compute weights for edges using overlapping heterozygous k-mers. Case (a) is a perfect case and the weight of edge (*x, x^r^*) is 2. Case (b), the left-side extension from *(TACCG,TACAG)* terminates when no left-overlapping pair of heterozygous k-mers is found and the right-side extension from (*CCGCA,CAGCA*) terminates when two pairs of right-overlapping heterozygous k-mers are found. Therefore, the weight of edge (*x, x*′) is 1.

More specifically, as for an isolated SNP k-mer pair, Kmer2SNP checks two properties this pair is supported by other overlapping heterozygous k-mers. A pair of k-mers (*y, y*′) is called *left-overlapping (right-overlapping)* with another pairs of k-mers (*x,x*′) if *x*[1,*k* – 1] = *y*[2, *k*], *x*′[1,*k* – 1] = *y*′[2,*k*] and *y*[1] = *y*′[1] (*x*[2, *k*] = *y*[1,*k* – 1], *x*′[2, *k*] = *y*′[1,*k* – 1] and *y*[*k*] = *y*′[*k*]). For example, (*AAGCA, AAGTA*) is left-overlapping with *(AGCAC,AGTAC*) in Fig. 3 (a). Kmer2SNP iteratively recruits left-overlapping and right-overlapping pairs of heterozygous k-mers to extend isolated SNP k-mer pairs on both sides. The extension on the left (right) side terminates if Kmer2SNP fails to find a unique pair of left-overlapping (right-overlapping) heterozygous k-mers which still covers the isolated SNP between. For example, the left-side extension from (*TACCG,TACAG*) terminates when no left-overlapping pair of heterozygous k-mers is found and the right-side extension from (*CCGCA,CAGCA*) terminates when two pairs of right-overlapping heterozygous k-mers are found in Fig. 3 (b). The *left (or right) extendable length* of an isolated SNP k-mer pair (*x, x*′), *l*(*x, x*′) (or *r*(*x, x*′)), is defined as the number of left-overlapping (right-overlapping) pairs of heterozygous k-mers recruited in the extension.

For example, *l*(*x,x*′) = *r*(*x,x*′) = 2 in Fig. 3 (a) and *l*(*x,x*′) = *r*(*x,x*′) = 1 in Fig. 3 (b). Finally, the weight of an edge between an isolated SNP k-mer pair (*x, x*′) is defined as min{*l*(*x,x*′), *r*(*x,x*′)}. Clearly, the edge weight between *x* and *x*′ in Fig. 3 (b) is 1. Please refer to the Supplementary Section 2 for the pseudocode to compute the weight of an edge between an isolated SNP k-mer pair (*x, x*′).

Similarly, we also introduce weights for edges between the corresponding nonisolated SNP k-mer pairs. We expect the True-Positive (TP) edges have higher weights than False-Positive (FP) edges, e.g., the mean weight of TP edges is 13.98 while the mean weight of FP edges is 1.19 in the same running example dataset HG-C_22_. In the next step, we thus use the maximum weight matching to select more confident SNPs.

#### Step 4 & 5 Compute maximum weight matching and output SNPs

According to the previous section, the more weight that Kmer2SNP assigns to an edge, the more likely the corresponding SNP is true. Therefore, Kmer2SNP computes the *maximum weight matching* in the above heterozygous k-mer graph, where the *maximum weight matching* is a set of pairwise non-adjacent edges in which the sum of weights is maximized. Kmer2SNP identifies all the connected components and uses python package NetworkX [11] to compute the *maximum weight matching* for each component with at least three vertices. This can be done efficiently thanks to the relatively small size of connected components in the graph.

The edges in the maximum weight matching are then converted back to SNPs as pairs of k-mer. Kmer2SNP further checks the weight distributions of edges in the maximum weight matching and automatically select a weight threshold to filter edges with lower weight. Please refer to the Supplementary Section 3 for more details. As each non-isolated SNP corresponds to two pairs of non-isolated SNP k-mers (refer to Fig. 1) and this non-isolated SNP is included in the output only if both edges are selected in the final maximum weight matching.

## 3 Results

### 3.1 Benchmarking Summary

**Datasets** We use the following datasets for our evaluations.

1. **HG-C**_*N*_. Simulated datasets are generate from Chromosome *N* using trio-phased haplotypes of individual NA12878 [10]. Illumina Hiseq reads with different coverages on different chromosomes are simulated by ART [12] Version 2.5.8.
2. **NA12878** and **NA24385**. NA12878 and NA24385 contain real 300X Illumina HiSeq reads aligned to different chromosomes of two individuals (NA12878 and NA24385) and are downloaded from NCBI [1, 2]. The trio-phased haplotypes for NA12878 [10] and NA24385 [3, 36, 37] are used as the ground-truth.
3. **Fungal**. The phased *Candida albicans SC5314* reference genome (version A22) is downloaded from Candida Genome Data [4]. The size of this fungal genome is 14.3 Mb and its heterozygous rate is 0.5% [14,23]. 100X Illumina Hiseq reads (50X for each haplotype) are simulated by ART [12].

#### Baseline Methods

We compare Kmer2SNP against two state-of-the-art reference-free SNP calling tools, DiscoSNP++ [33, 26] and EBWT2SNP [28], as they show the best results in recent benchmarking [26, 28]. Please refer to the Supplementary Section 6 for detailed parameter setting. As for hybrid approaches, we use SPAdes [6] and SGA [30] to assemble Illumina reads into contigs. We further use BWA [16] to align reads to the assembled contigs and apply two popular SNP calling pipelines, SAMtools (bcftools) [24] and GATK [5], to call SNPs. These *hybrid* approaches are called SPAdes+GATK, SGA+SAMtools, etc.

#### Ground truth and Comparison Metrics

Note reference-free approaches cannot assign SNPs to known genomic positions as the reference genome is unknown. For NA12878 and NA24385, we download the reference genomes along with SNP annotations from [10, 3, 36, 37] to generate heterozygous k-mer pairs as ground-truth. For Fungal, we download two homologous sets of chromosomes (download from [4]), and then use BLASR [7] to align them to obtain heterozygous k-mer pairs.

As different reference-free approaches for SNP-calling may have different output formats, we convert all output SNPs into heterozygous k-mer pairs to make a fair comparison. By default, k is set to be 31 such that at least 98% of heterozygous k-mers are able to assigned to unique genomic positions when the reference genome is known in the above datasets. Three metrics, recall, precision and F1-score, are used to evaluate the performance of SNP calling. We also record the CPU time and refer to the Supplementary Section 6 for the running environment.

### 3.2 Results on HG-C_*N*_ datasets

Similar to the experiments conducted DiscoSNP++ [33, 26] and EBWT2SNP [28], we perform extensive experiments by varying the k-mer sizes and the sequencing coverages on different chromosomes. Among the many combinations of parameters tested, we show some representative results on HG-C_16_ and HG-C_22_. HG-C_16_ consists of 80X simulated Illumina Hiseq reads (40X for each haplotype) of Chromosome 16 while HG-C_22_ consists of simulated Illumina Hiseq reads with different coverages of Chromosome 22.

#### The choice of k-mer sizes

The k-mer size is an important parameter for Kmer2SNP, DiscoSNP++ and EBWT2SNP. DiscoSNP++ shows that the k-mer size has a limited impact on the SNP calling quality [33, 26] and performs experiments by setting k=31. EBWT2SNP [28] chooses a sufficiently large k-mer (k=31 in all the experiments) such that a k-mer is expected to appear at most once in the genome. KMER2SNP also achieves stable performance across different k-mer sizes (refer to Figure S2 in the Supplementary Section 4). Therefore, in the following experiments, the k-mer size is chosen to be 31 by default.

#### The affect of different coverages

Fig. 4 shows that Kmer2SNP outperforms DiscoSNP++ and EBWT2SNP on recall, precision and F1-score while being an order of magnitude faster for SNP calling on different coverage of HG-*C*_22_. Moreover, with the increase of read coverages, the running time of DiscoSNP++ and EBWT2SNP increases significantly while KMER2SNP still maintains low running time.

**Fig. 4.**
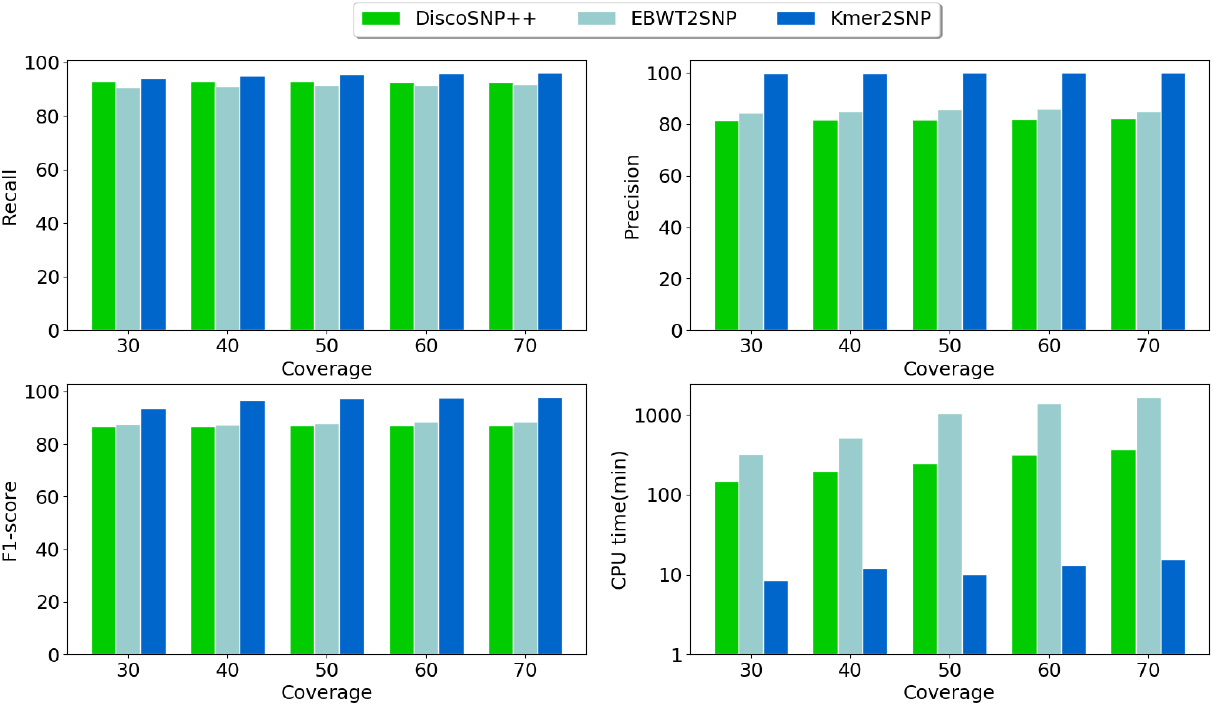
SNP calling on different read coverages of Human Chromosome 22 (HG-C_22_).

### 3.3 Results on NA12878 and NA24385 datasets

We benchmark Kmer2SNP against DiscoSNP++, EBWT2SNP and hybrid approaches on real 300X Illumina HiSeq reads of Chromosome 22 of NA12878 and NA24385. The SNP calling results on NA12878 (Chromosome 22) and NA24385 (Chromosome 22) are shown in Table 1 and Table S1 in Supplementary Section 5. Both tables demonstrate that Kmer2SNP outperforms existing reference-free SNP calling tools in both quality and scalability. Consistent with previous results [33], although hybrid approaches achieve high recall rates, they suffer seriously from errors in assemblies of complex genomes, thus call many false positive SNPs and result in low precision rates.

**Table 1.** Performance of SNP calling on NA12878 (Chromosome 22) dataset [1].

| Type | Tools | Recall | Precision | F1-score | CPU time (h) |
| --- | --- | --- | --- | --- | --- |
| Hybrid<br>(assembly-based) | SPAdes+GATK | **94.48** | 43.40 | 59.48 | 80.53 |
|  | SPAdes+Samtools | 94.13 | 46.96 | 62.66 | 64.98 |
|  | SGA+GATK | 77.74 | 44.19 | 56.35 | 57.31 |
|  | SGA+Samtools | 77.25 | 43.68 | 55.81 | 41.26 |
| Assembly-free | DiscoSNP++ | 76.99 | 76.36 | 76.67 | 8.88 |
|  | EBWT2SNP | 85.34 | 56.17 | 67.75 | 47.16 |
|  | Kmer2SNP | 83.60 | **84.03** | **83.81** | **0.46** |

### 3.4 Results on Fungal dataset

Table 2 summarizes the SNP calling results of Kmer2SNP, DiscoSNP++ and EBWT2SNP as well as hybrid approaches for SNP calling on *Candida albicans SC5314* dataset (100X simulated Illumina reads, 50X for each haplotype). Similar to the above experiments results, Kmer2SNP outperforms existing reference-free approaches including DiscoSNP++, EBWT2SNP and hybrid approaches. Note the precision of hybrid approaches improves as this genome does not contain complex repeat structures as in the human genome, however, the overall performance of hybrid approaches is still not as good as Kmer2SNP.

**Table 2.** Performance of SNP calling on the Fungal dataset (100X Illumina Hiseq reads from the phased *Candida albicans SC5314* reference genome (version A22) [4]).

| Type | Tools | Recall | Precision | F1-Score | CPU time (h) |
| --- | --- | --- | --- | --- | --- |
| Hybrid<br>(Assembly-based) | SPAdes+GATK | 77.42 | 91.80 | 84.00 | 15.32 |
|  | SPAdes+SAMtools | 76.87 | 92.18 | 83.83 | 10.09 |
|  | SGA+GATK | 62.52 | 94.00 | 75.09 | 12.15 |
|  | SGA+SAMtools | 62.50 | 93.01 | 74.76 | 6.92 |
| Assembly-free | DiscoSNP++ | 76.29 | 96.88 | 85.36 | 2.41 |
|  | EBWT2SNP | 68.55 | 97.51 | 80.50 | 4.91 |
|  | Kmer2SNP | **78.09** | **97.57** | **86.75** | **0.11** |

## 4 Conclusion and Discussion

Kmer2SNP shows the potential of calling SNPs directly from raw reads when the reference genome is not available. Kmer2SNP introduces a graph model on k-mers and the SNP calling problem is to find the maximum weight matching in the graph. Kmer2SNP outperforms other reference-free approaches in SNP calling quality while being an order of magnitude faster.

Kmer2SNP has several limitations. The current implementation of Kmer2SNP only supports at most two non-isolated SNPs in a heterozygous k-mer pair for efficiency purposes. How to extend Kmer2SNP to find more non-isolated SNPs is worth investigating. Currently, Kmer2SNP is only applicable to diploid genomes and its potential for handling polyploid genomes needs more analysis. Last but not least, Kmer2SNP may benefit from varying the k-mer sizes in building the vertex set (similar to the work in [20]) and introducing a probabilistic model in computing the maximum weight matching (similar to the one in [25]).

## Footnotes

1 Note that k-mers from paralogous sequence variants (PSVs) belong to homozygous k-mers and thus are filtered.

