## Supplementary Material for "Kmer2SNP: reference-free SNP calling from raw reads based on matching"

Yanbo Li[1], and Yu Lin[1,*]

#### 1 Construct $h2$-edges

According to the definition of non-isolated k-mer pairs, there is an $h2$-edge between two k-mers $x$ and $x'$ if and only if $x[(k+1)/2] \neq x'[(k+1)/2]$ and $h(x, x') = 2$, where k is odd and h() computes the hamming distance. For each k-mer $x$, Kmer2SNP uses the prefix $x[1, (k-1)/2]$ and suffix $x[(k-1)/2, k]$ respectively as keys to compute its index in hash table. Two k-mers $x$ and $x'$ are connected by an h2-edge only if they have the same index. Note that Kmer2SNP needs to verify the hamming distances between k-mers with the same index to add h2-edges.

#### 2 Pseudocode to compute the weight of an edge $(x, x')$ in the heterozygous k-mer graph

**Algorithm 1:** Compute the weight of an edge $(x, x')$ in the heterozygous k-mer graph

**Input:** an edge between an isolated SNP k-mer pair $(x, x')$, the heterozygous k-mer graph
**Output:** the weight of this edge $(x, x')$

$l_{temp} \leftarrow 0, r_{temp} \leftarrow 0$ ;
$x_{current} \leftarrow x, x'_{current} \leftarrow x'$;
**while** *there exists a unique heterozygous k-mer pair (y, y') left-overlapping with* $(x_{current}, x'_{current})$ **do**
    $l_{temp}{+}{+}$;
    $(x_{current}, x'_{current}) \leftarrow (y, y')$;
    **if** $l_{temp} > k/2 - 1$ **then**
        *break;*
    **end**
**end**
$x_{current} \leftarrow x, x'_{current} \leftarrow x'$;
**while** *there exists a unique heterozygous k-mer pair (y, y') right-overlapping with* $(x_{current}, x'_{current})$ **do**
    $r_{temp}{+}{+}$;
    $(x_{current}, x'_{current}) \leftarrow (y, y')$;
    **if** $l_{temp} > k/2 - 1$ **then**
        *break;*
    **end**
**end**
**return** $min\{l_{temp}, r_{temp}\}$;

#### 3 Weight distribution of edges in the maximum weight matching and a filtering threshold in the postprocessing

Figure S1 shows the weight distribution of edges in the maximum weight matching for HG-$C_{22}$ (30X coverage on each haplotype). A filtering threshold is selected as the first local minimum in the discrete approximation of the first order derivative of the weight distribution. More specifically, let $W(i)$ be the number of edges of weight $i$ and $\Delta_i = W(i) - W(i+1)$. The filtering threshold is selected as the smallest $i$ such that $\Delta(i) < \Delta(i+1)$ and $\Delta(i) < \Delta(i-1)$. In Figure S1, Kmer2SNP automatically selects a filtering threshold as 3 and thus Kmer2SNP will filter edges whose weights are less than 3.

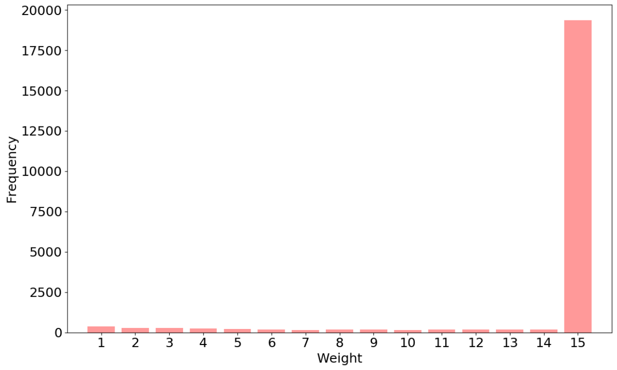


Figure S1: The weight distribution of edges in the maximum weight matching for HG-$C_{22}$ (30X coverage on each haplotype) dataset

### 4 The choice of k-mer sizes

The k-mer size is an important parameter for Kmer2SNP, DiscoSNP++ and EBWT2SNP.

- DiscoSNP++ shows that the k-mer size has a limited impact on the variant calling quality uricaru2014reference,peterlongo2017discosnp++ and performs experiments by setting k to be 31.
- EBWT2SNP prezza2018detecting,prezza2019snps chooses a sufficiently large k-mer (k=31 in all the experiments) such that a k-mer is expected to appear at most once in the genome.
- Figure S2 shows that KMER2SNP also achieves stable performance across different k-mer sizes.

Therefore, in the following experiments, the k-mer size is chosen to be 31 by default.

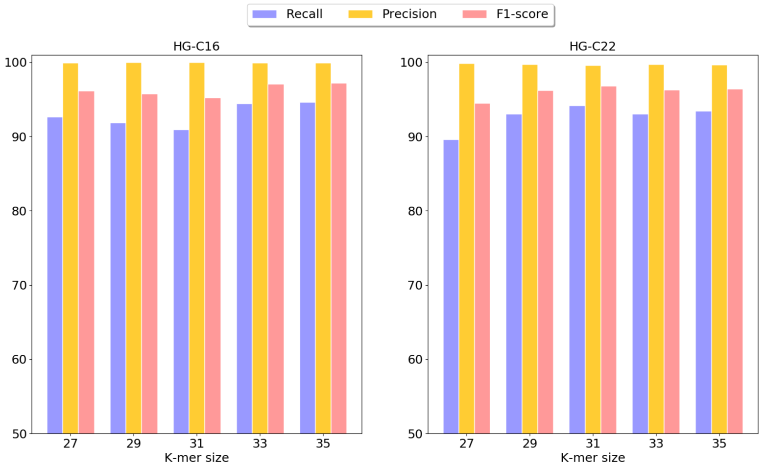


Figure S2: Different choices of the k-mer size for KMER2SNP (recall, precision, F1 and) on HG-$C_{16}$ (40X coverage on each haplotype) and HG-$C_{22}$ (30X coverage on each haplotype).

### 5 The results on NA24385 (Chromosome 22)

Table S1 again demonstrates that Kmer2SNP outperforms existing reference-free SNP calling tools in both quality and scalability. Note that although hybrid approaches achieves high recall rates, their precesion rates are very low, probably due to errors in assemblies.

Table S1: Performance of SNP calling of NA24385 (Chromosome 22) dataset NA24385

| Type | Tools | Recall | Precision | F1-score | CPU time (h) |
|---|---|---|---|---|---|
| Hybrid(assembly-based) | SPAdes+GATK | **94.87** | 30.30 | 45.93 | 84.6 |
| | SPAdes+SAMtools | 94.53 | 35.52 | 51.64 | 67.27 |
| | SGA+GATK | 76.45 | 37.14 | 49.99 | 67.66 |
| | SGA+SAMtools | 75.63 | 37.31 | 49.97 | 48.5 |
| Assembly-free | DiscoSNP++ | 65.48 | 61.89 | 63.63 | 9.40 |
| | EBWT2SNP | 83.29 | 49.84 | 62.36 | 41.75 |
| | Kmer2SNP | 78.23 | **66.24** | **71.74** | **0.44** |

### 6 Running environment and commands

We run all those software on a server with 1000GB of RAM and Intel(R) Xeon(R) CPU E7- 4870 @ 2.40GHz * 80 CPU processor.

- EBWT2SNP
  /path/EBWT-v2/egap/eGap -m 4096 chr22.2strand.fasta
  /path/EBWT-v2/ebwt2snp-v2/build/ebwt2snp -1 chr22.2strand.fasta.bwt -o output.snp -t 0
  /path/EBWT-v2/ebwt2snp-v2/build/filter_snp output.snp low_cov high_cov > output.filter.snp
  (Note that two parameters, low_cov and high_cov, are decided by `DSK` and `findGSE`, same as the k-mer frequence range used in Kmer2SNP)

- DiscoSNP
  /path/DiscoSnp/run_discoSnp++.sh -r fof.txt -T -u 10

- Alignment for NGS
  bwa index one_haplotypes.fasta
  bwa mem -M -t 30 -R '@RG\tID:foo\tSM:bar' one_haplotypes.fasta single_dat.fq > one_haplotype_single.sam
  samtools view -@ 20 -S -bF 4 -q 1
  one_haplotype_single.sam > filtered.bam
  samtool sort -@ 20 filtered.bam -o filtered.sorted.bam
  samtools index filtered.sorted.bam

- SAMtools calling
  samtools faidx one_haplotypes.fasta
  bcftools mpileup -f one_haplotypes.fasta one_haplotype_single_sorted.bam | bcftools call -mv -Ob -o calls.bcf
  bcftools view calls.bcf | vcfutils.pl varFilter - > calls.vcf

- GATK calling
  gatk SortSam -I filtered.sorted.bam -O gatk.sort.bam -SO coordinate
  gatk MarkDuplicates -I gatk.sort.bam -O gatk.dd.bam --METRICS_FILE gatk.dd.metrics --MAX_FILE_HANDLES_FOR_READ_ENDS_MAP 1000
  samtools faidx one_haplotypes.fasta
  gatk CreateSequenceDictionary -R one_haplotypes.fasta -O one_haplotypes.dict
  samtools index gatk.dd.bam
  gatk HaplotypeCaller -R one_haplotypes.fasta -I gatk.dd.bam -O raw.snp.indels.vcf
